# Migration of di-(2-ethylhexyl) phthalate from the plastic film into the air and vegetables in the greenhouses

**DOI:** 10.1101/479469

**Authors:** Xiaowei Fu, Bin Huang

## Abstract

The aim of this study was to investigate di-2-ethylhexyl phthalate (DEHP) from plastic film of the greenhouse to the air and the vegetables grown in the greenhouse. This paper presents the migration of di-2-ethylhexyl phthalate (DEHP) by determining the di-2-ethylhexyl phthalate (DEHP) concentration in air and vegetables in greenhouses with the change of temperature in a day and a half year. The determination of di-2-ethylhexyl phthalate (DEHP) content in the air and the vegetables was performed by gas chromatography-mass spectrometry. The results showed that di-2-ethylhexyl phthalate (DEHP) content in the greenhouse air increased with increasing temperature during one day, but di-2-ethylhexyl phthalate (DEHP) content changes in vegetable were different from those in air, and which depends on the exposure time. For the monthly content changes, di-2-ethylhexyl phthalate (DEHP) content in air first increased to a maximum and then decreased with increasing time and temperature. The di-2-ethylhexyl phthalate (DEHP) content changes of six vegetables not only were related with di-2-ethylhexyl phthalate (DEHP) content in air and temperature, but also with varieties of vegetables.

## Introduction

Plastic greenhouses are a form of horticulture which plays a special role for supplying off-season vegetables. As food-safety is increasingly concerned, the safety evaluation of vegetable is not limited to heavy metals and pesticide residues, but also organic contaminants [9]. Contamination of vegetables by the plasticizers is being received serious attention. The most used plasticizer, di-(2-ethylhexyl)phthalate (DEHP) is easily released to environment from plastic products due to its intability, fluidity and volatility[1] since DEHP is not chemically bonded to the polymer of plastics. Therefore, DEHP from the greenhouse film is possible of an organic contaminant of vegetables cultivated in greenhouses.

Migration of DEHP from the plastic products to several foods was reported. Zhang et al.[19] reported that meat wrapped with PVC film could be contaminated by DEHP, and the migrated amount of DEHP depended on the temperature and time that the meat wrapped by plastic film. Goulas[7] studied that the migration of DEHP from food-grade polyvinyl chloride film into hard and soft cheeses, and the result showed that the longer the exposure, the higher levels of DEHP in cheese. The uptake of DEHP from plastic mulch film used in field cultivation by 10 vegetable plants was investigated by experiments carried out in pots by Du et al. [5,6]. The results showed that DEHP was transferred from the mulch film to all vegetable plants, and *Benincasa hispida*, *Cucumis sativu*, *Cucurbita moschata* and *Brassica parachinensis* could accumulate high level of DEHP from mulch plastic film giving dietary exposure over or close to the daily intake limit for DEHP. The air DEHP from the plastic production building could cause contamination of vegetables according to a determination of DEHP concentrations in the atmosphere and four vegetable crops cultivated on land surrounding a plastic production factory [6]. However, no data is available concerning the migration of DEHP from the plastic film into vegetables in the greenhouses. This paper presents the migration of DEHP by determining the DEHP concentration in air and vegetables in greenhouses with the change of temperature in a day and a half year.

## Materials and methods

### Solvents and reagents

All solvents and reagents used for extraction were of analytical grade and purchased from Huadong Chemicals Company (Hangzhou, China). DEHP and chromatographically pure chloroform for dissolving DEHP standard was purchased from Sigma Shanghai Division (Shanghai, China).

### Film material

The film was manufactured by Liaochen Plastic Film (Shangdong, China) and was used for the field-plant experiments.

### Plants used for experiments

Some vegetables cultivated in the plastic-film greenhouse including bok choy (*B. chinensis*), celery (*A. graveolens*), spinach (*S. oleracea L.*), cabbage (*B. oleracea*), lettuce (*L. sativa*), edible amaranth (*A. mangostanus*), were selected for testing DEHP uptake.

### Sampling

#### The daily variation of DEHP from the greenhouse films

The edible parts of bok choy (*B. chinensis*) and the air samples in the greenhouses were collected at the same time for DEHP determination. The air samples were collected at 5:00, 8:00, 11:00, 14:00, 16:00, 19:00 and 22:00 in 3 consecutive days by the activated carbon adsorption method[17,18]. And the temperature at each time was recorded. For each DEHP determination, three replications were carried out.

#### The monthly variation of DEHP from the greenhouse films

From December 1, 2010 to June 10, 2011, the air samples and six vegetables were collected in the tenth day of each month at 9:00 am. Six vegetables were planted in the first day of each month and sampled in the 10^th^ day of next month. The air in the greenhouse was sampled by the activated carbon adsorption method, and the temperature at each time was recorded. For each DEHP determination, three replications were carried out.

#### Determination of DEHP

Extraction of DEHP

Vegetable samples were freeze-dried, ground and sieved to less than 0.2 mm. For each sample, 5-10 g was extracted in a Soxhlet extractor for 24 h with 200 mL acetone/dichloromethane (1:1,v/v) in a water-bath at 75 ℃ . The extracts were reduced to 5.0 mL using a rotary evaporator in a water-bath at 50 ℃ . The concentrated sample was subjected to clean-up.

Air samples: For each sample, 10 g was extracted in a Soxhlet extractor for 24 h with 200 mL acetone/dichloromethane (1:1,v/v) in a water-bath at 55 ℃ . The extracts were reduced to 5.0 ml using a rotary evaporator in a water-bath at 50 ℃ . The concentrated sample was subjected to clean-up.

Clean-up

The concentrated samples were loaded on a combined column of silica gel and alumina. The glass chromatography column 25 cm long, 1c m I.D.), was packed with 3 cm alumina plus 10 cm silica, followed by 2 cm anhydrous sodium sulfate. Dichloromethane (20 mL) was used for elution. The collected fraction was reduced to 0.5 mL under a gentle stream of nitrogen.

### Gas chromatography-mass spectrometry

The cleaned samples were analyzed by GC-MS [8,13] using a Hewlett-Packard 5890/5971 GC-MSD (Agilent Technologies, Palo Alto, CA, USA) equipped with an HP-5 trace analysis column (30 m, 0.32 mm I.D., 0.25 mm film thickness). The GC oven temperature was held at 150 ℃ for 3 min, and programmed to increase at 20℃ min^−1^ to 300 ℃ and finally held at 300 ℃ for 3 min.

The temperature of the injector was 250 °C. Helium was the carrier gas at a linear flow-rate of 20.7 cm s^−1^. Full scan electron ionization data was obtained as follows: solvent delay 5 min, electron ionization energy 70 eV, source temperature 200 °C, emission current 150 μA, scan rate 4 scan s^−1^, detector voltage 350 V. The DEHP level in the sample was taken as the average of three injections. The amounts of DEHP were calculated from a calibration curve: y=0.0014 x + 0.0006 (r^2^=0.9911) for air samples and y=0.0008 x + 0.0003 (r^2^=0.9848) for vegetable samples. The final contents of the vegetables were expressed as μg g^−1^ based on the amounts of the dried samples.

## Results and discussion

### The daily variation of DEHP from the greenhouse films The daily variation of DEHP in the greenhouse air

The concentration of DEHP in the greenhouse air was determined by gas chromatography-mass spectrometry. The results show that the DEHP content in air increased with the increase of the greenhouse temperature (Fig.1). The air DEHP content was the lowest in 5:00, and then as the temperature rose, the content was gradually increased. To 2 pm when the temperature was the highest point, the DEHP content reached also the highest value. Then as the temperature decreased, the content was slowly declining. It was evident that the DEHP content in the greenhouse air was closely related with the greenhouse temperature.

During the day as temperatures rise, the temperature in the greenhouse also will increase. And it also accelerated the release rate of DEHP from the film. That is, DEHP content in air increased, and reached the highest point when the temperature reaches the maximum.

As the day time, the temperature gradually decreased. Subsequently the released rate of DEHP from the greenhouse films was affected, which was bound to affect the release of DEHP. Coupled with the greenhouse was not a closed container, or DEHP may also be absorbed by plants or soil in the greenhouse. Therefore, the temperature dropped, the DEHP content in air also declined.

**Figure 1.**
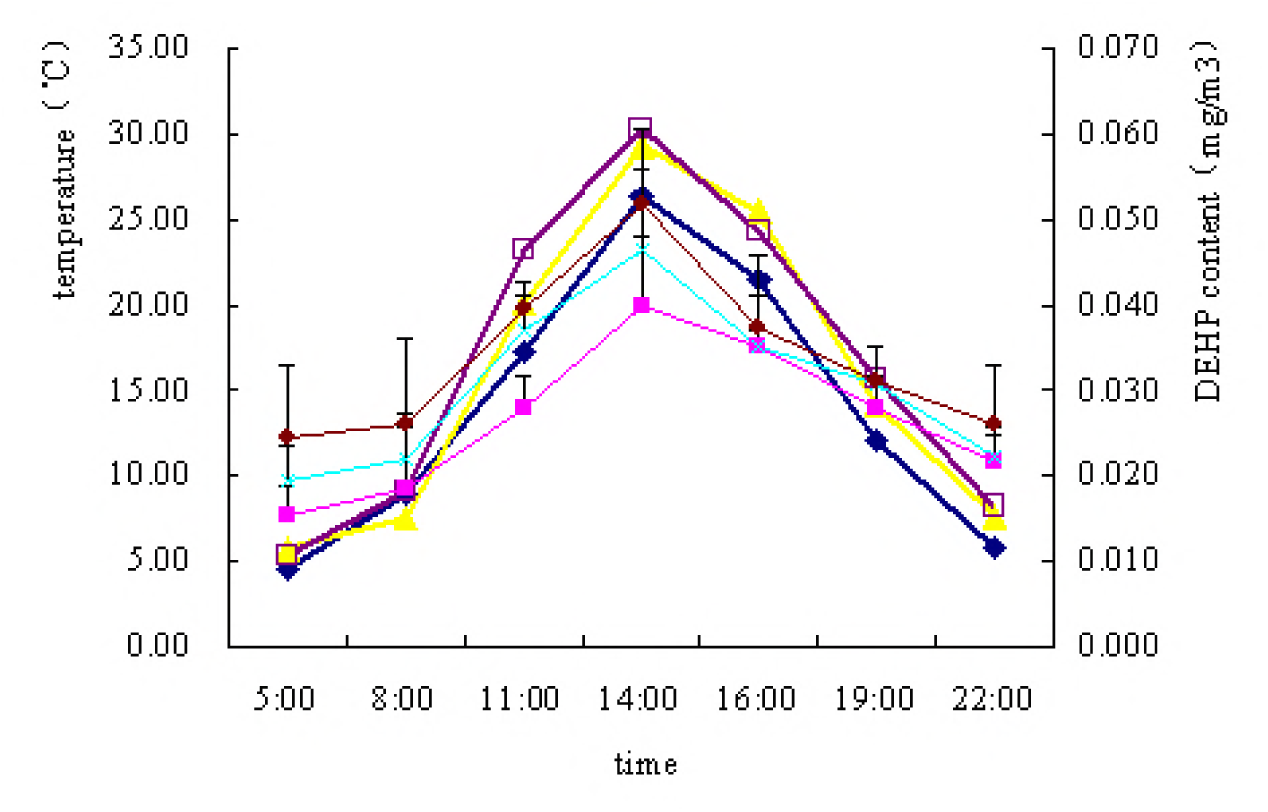
The change of temperature and DEHP content of air in the greenhouse during a day. March 24, 2011; ◆temperature; ■DEHP content March 25, 2011; ▲temperature; ×DEHP content March 26, 2011; □temperature; ○DEHP content

Wang et al.[16] reported that the characteristics of distribution of Phthalate esters (PAEs) in vegetable greenhouses. Our results are in agreement with their observations.

### The daily variation of DEHP in the greenhouse vegetables

Fig. 2 show that the DEHP content in the greenhouse vegetable was increased from morning to night. For the March 24, March 25, and March 26, 2011 samples, DEHP content were slowly increasing as time went on.

**Figure 2.**
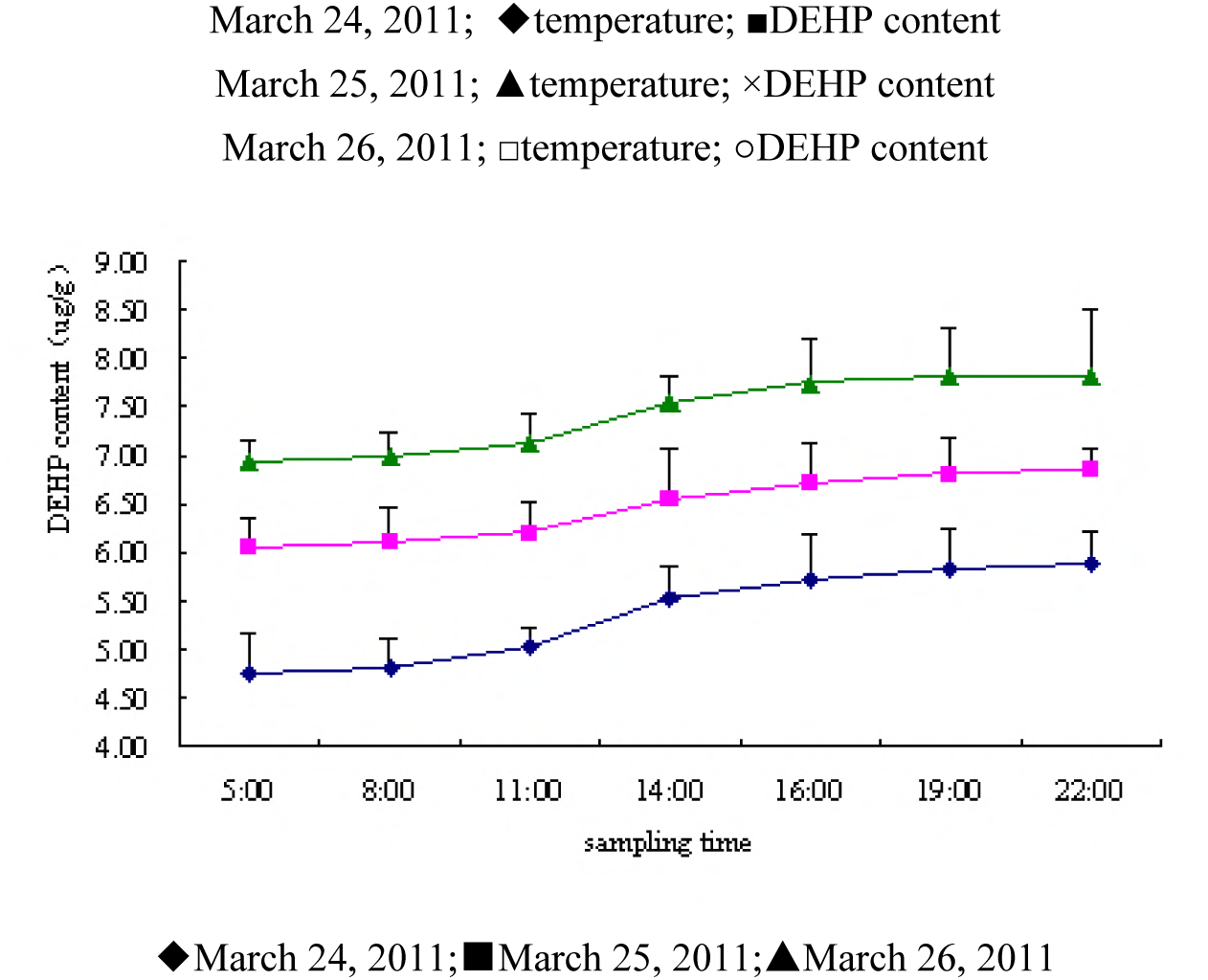
The change of DEHP content of *B.chinensis* cultivated in the greenhouse.

DEHP daily changes in vegetables were not consistent with those in the air (Fig.3). DEHP daily content in air increased and then decreased, while the daily content in vegetables was increasing from the morning to the evening.

**Figure 3.**
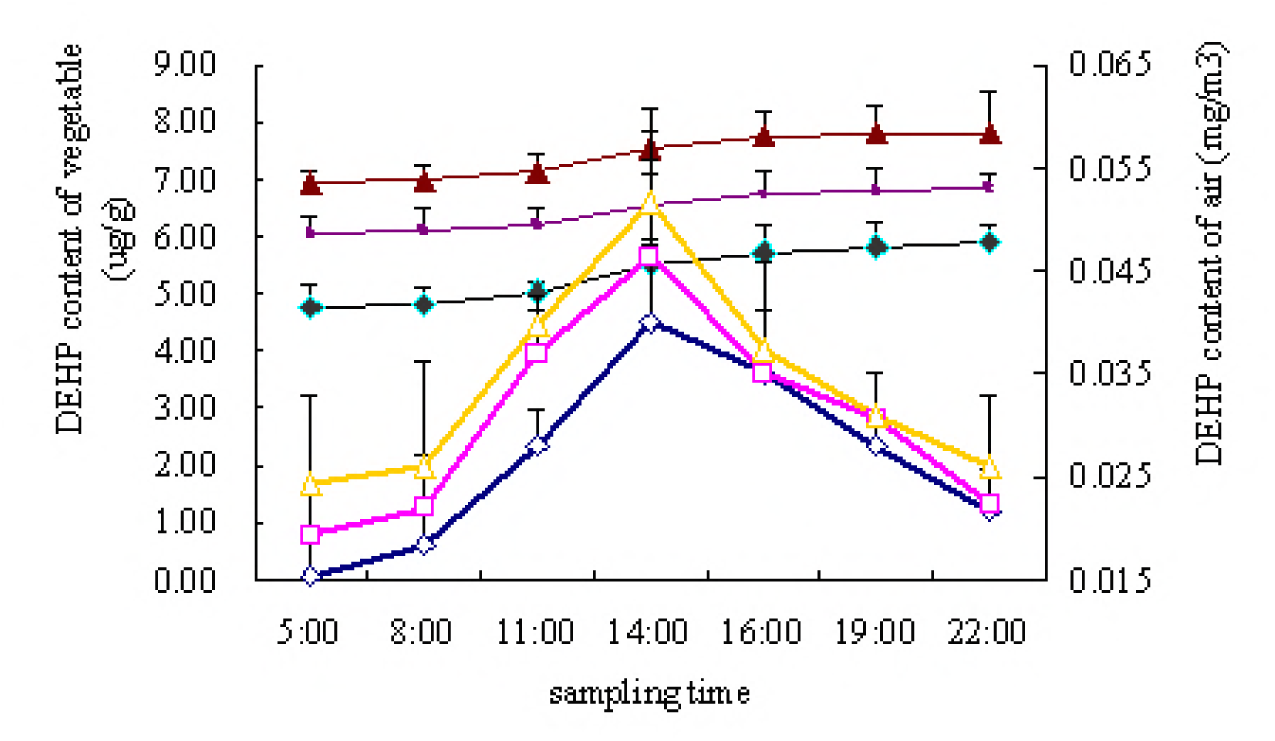
The comparison between the changes of DEHP content of air and *B.chinensis* in the greenhouse. March 24, 2011; ◇air; ◆*B.chinensis* March 25, 2011; □air; ■*B.chinensis* March 26, 2011; △air; ▲*B.chinensis*

When the DEHP in the air was the highest levels, it does not contain the maximum amount of DEHP in vegetables. This can be inferred that the DEHP concentration in vegetables was related not only with the content of DEHP in the air, but also the time of absorption.

### The monthly variation of DEHP from the greenhouse films The monthly variation of DEHP in the greenhouse air

With the increase in the greenhouse age, levels of DEHP in the air firstly slowly rose and then rapidly rose and slowly declined followed two months (Fig.4). As time went on, the total amount of DEHP in films was gradually reduced, so that their emission was also reduced. Therefore, the concentration of DEHP in the air gradually decreased after reaching a certain concentration.

**Figure 4.**
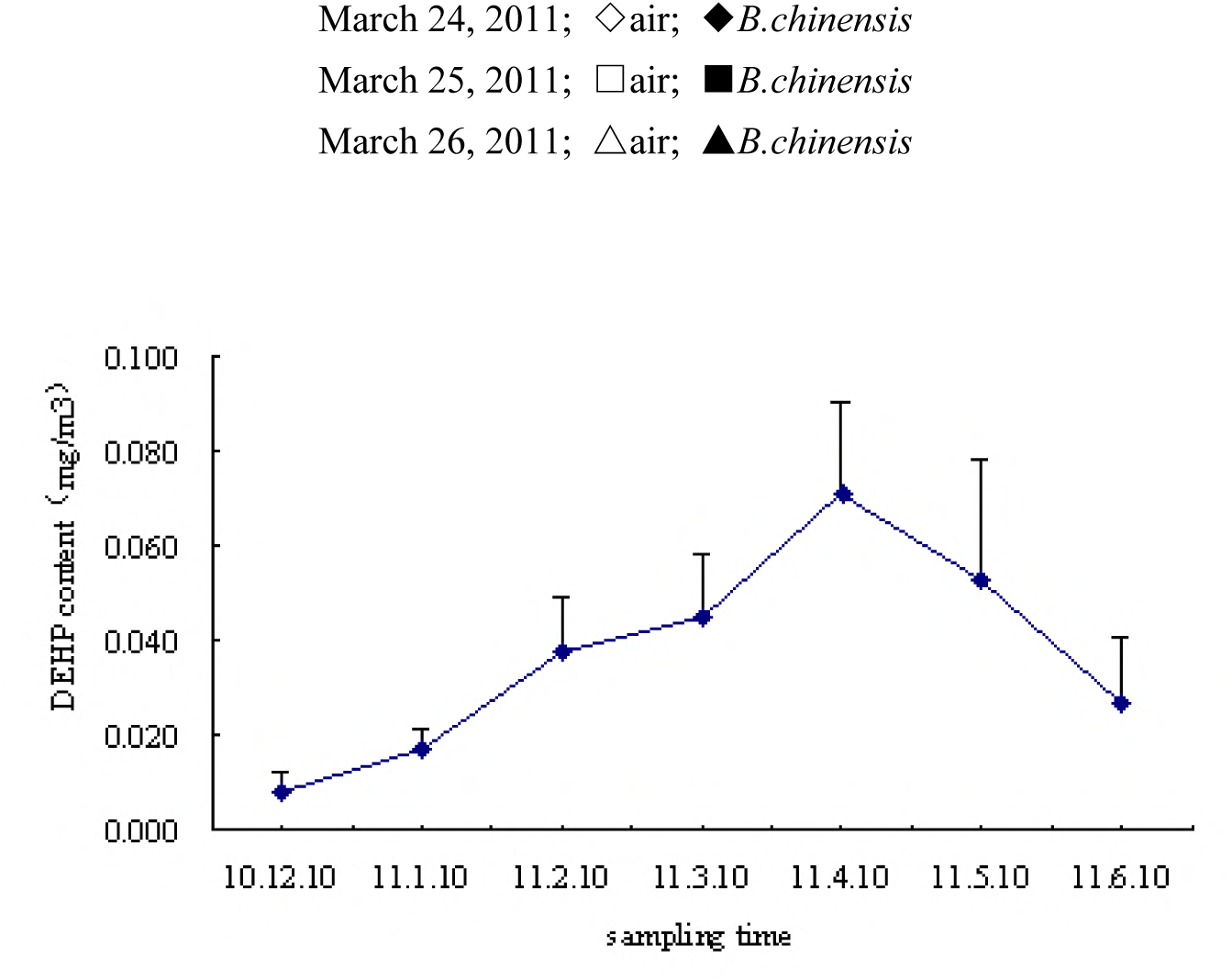
The monthly variation of DEHP content of air in the greenhouse.

DEHP content in the air was also associated with the temperature. The higher the temperature, the faster the molecular thermal motion in films, which helps to evaporation and leaching of DEHP from the films. From December 10, 2010 to April 10, 2011, with the gradual increase in ambient temperature, which will affect the micro-climate in the greenhouse, the concentration of DEHP of the air in the greenhouse increased slowly to reach maximum concentration in April. With levels of DEHP of the films decreasing, which will reduce its total dissolution, and thus the concentration of greenhouse air was gradually decreased (Fig.4). During May and June, although the temperature is relatively high, but the total DEHP content of films gradually reduced, which also reduced the emission of DEHP, and the number of open-air greenhouse were added, so levels of DEHP in air decreased. Zheng et al. [23] reported the migration of the PAEs from the PVC plastic products in the water environment. Wang et al.[16] studied the variations of the PAEs levels from winter to spring. These results are similar with our findings.

### The monthly variation of DEHP in the greenhouse vegetables

Fig.5 show that the content changes of DEHP in *L. sativa, S. oleracea L. and A. mangostanus* were somewhat regular. With increasing age of the greenhouse, the DEHP content in the three vegetables also increased before May 10, 2011. However, the levels of DEHP in the six vegetables had decreased in varying degrees from May 10, 2011 to June 10, 2011. This result is consistent with the variation of DEHP levels in the greenhouse air. There are two reasons to explain this result. One is that the total DEHP of the membrane decreased; second is that the greenhouse are required to increase ventilation times because of the high temperature.

**Figure 5.**
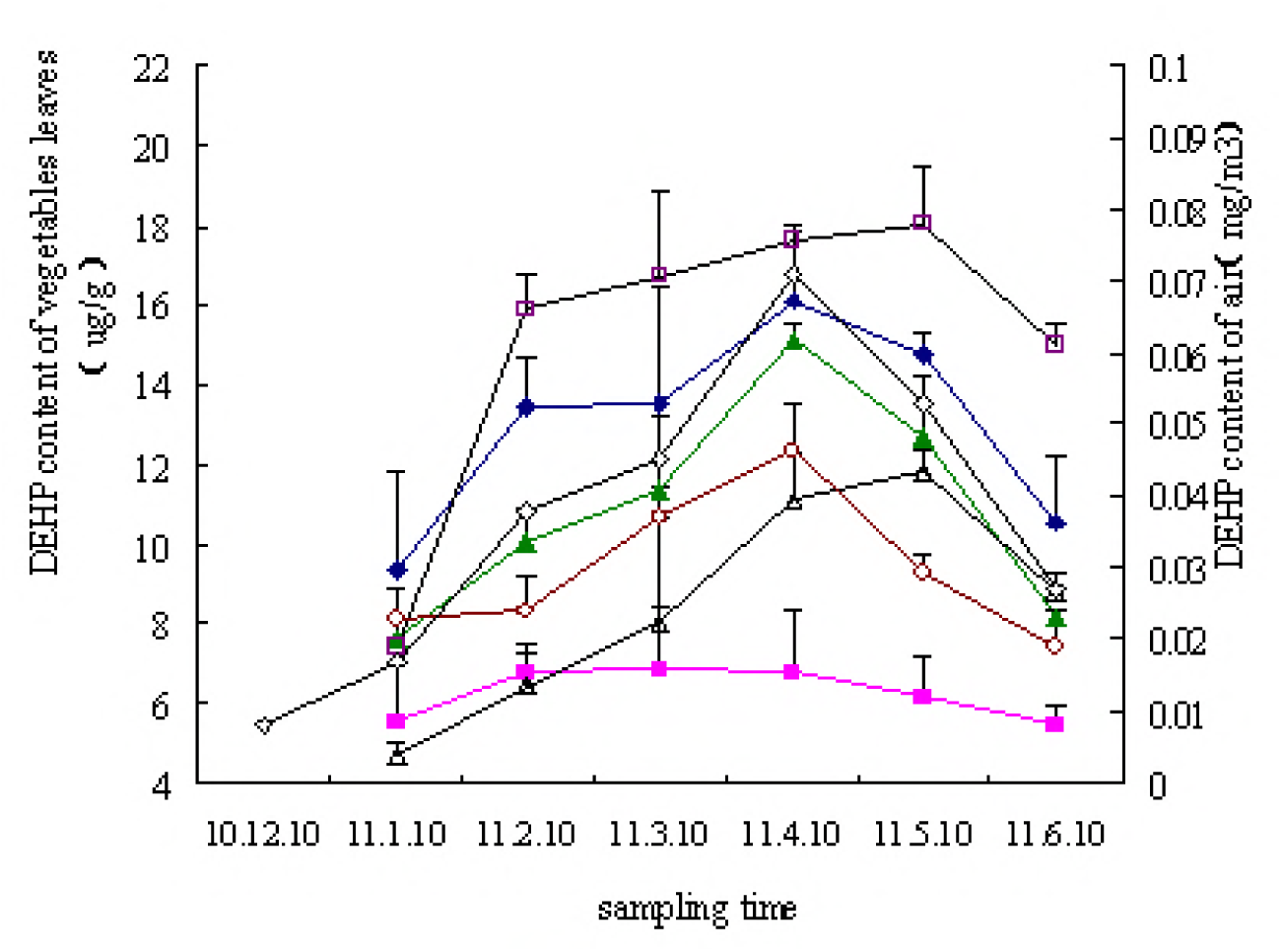
The monthly variation of DEHP content of the different vegetables from the greenhouse. (◆*B. chinensis*; ■*B. oleracea*; ▲*S. oleracea*; △*A. graveolens*; □*L. sativa*; ○*A. mangostanus*; ◇air)

Among the 6 vegetables, three kinds of vegetables *(L. sativa, B. chinensis and A*. *graveolens*) can be strong to absorb DEHP in air, followed by the other two vegetables (*S. oleracea L.and A. mangostanus*). While a vegetable (*B. oleracea*) is the weakest ability to absorb DEHP. The results concluded that the DEHP content in vegetables is related with the vegetable varieties. And different vegetables have different absorption or adsorption ability of DEHP. Wang Jiawen et al. [17] reported the DEHP pollution of vegetable from farmland and differences between vegetables species. contaminated by DEHP. Wang et al.[16]studied that the distribution of Phthalate esters in the greenhouse vegetables in the different ages. In this regard, our results are basically the same with their.

## Acknowledgements

This research was supported by a fund (LY16D010003) from the Zhejiang Provincial National Science Foundation of China.

## Author contributions statements

Xiaowei Fu wrote the main manuscript text and Bin Huang prepared figures 1–3. All authors reviewed the manuscript.

## Competing financial interests

The authors declare no competing financial interests.

